# A niche construction-based perspective on chemical evolution before replication and reproduction

**DOI:** 10.1101/2025.07.29.667458

**Authors:** François Papale, Kulish Yuri, Joseph P Bielawski, Louis-Patrick Haraoui

## Abstract

Conceptual analysis has, in the past decades, established that evolution by natural selection (ENS) can occur without reproduction (1–6). This theoretical advancement has significant imports for research on the origins of life that have yet to be explored. In this paper, we introduce and defend a niche construction-based account of evolution by natural selection without reproduction (ENSwR, pronounced like “answer”), which leads to three insights regarding Darwinian evolution at the origins of life. First, we show through ENSwR the plausibility of Darwinian evolution in the prebiotic world; second, ENSwR provides a novel explanatory framework that supports origins of life theories in which autocatalytic sets of chemical reactions play a central role; third, we argue that inorganic catalytic molecules (rather than autocatalytic sets (7–9)) are relevant units of selection to understand the origins of life prior to reproduction and replication. This third argument is especially important for the field, as it helps bridge the gap between chemical and biological evolution (10).

## 1. Introduction: evolution beyond replicators

Self-replication and reproduction are generally assumed to be necessary for evolution by natural selection (ENS) to occur (11–14). The centrality of this condition is such that self-replication is also believed by many to be a defining criterion of life (15, 16). Recent conceptual analysis, however, has shown that if ENS is defined as change in the distribution of variation across time that is due to fitness-affecting environmental interactions (i.e., to natural selection), then populations can undergo ENS even if their components, the units of selection, fail to reproduce or self-replicate (1–6). This insight has the potential to drastically alter the way we think about Darwinian evolution. It seems especially promising when it comes to understanding the evolutionary origins of the first reproducing entities, whether they be RNA, proteins or autocatalytic sets of chemical reactions (ACSs). In this paper, we show how the theory of evolution by natural selection *without reproduction* (ENSwR, pronounced like “answer”) can fill the gap between chemical and biological evolution, thereby expanding the explanatory reach of the Darwinian framework prior to and beyond biology in an innovative way. The framework we describe also strengthens origins of life theories in which metabolism and ACSs play a central role.

The body of theories regarding the origins of life have mostly been divided into two broad categories (17, 18): genetic-first approaches (e.g., RNA world (19, 20)) and metabolic-first approaches (e.g., autocatalytic networks (7)). Some have also argued for more pluralist models combining independent and convergent pathways (21). Despite important differences between these approaches to the origins of life, they share the belief that ENS is an important process for explaining the emergence of life (especially of the first cellular organisms) *and* that reproduction is necessary to get the process of ENS going (22). This leaves these approaches, in their current state at least, in a dire situation: if ENS requires reproduction of some sort, this means that the first reproducers/replicators cannot have resulted from a process of ENS. How, then, did they evolve?

In this paper, we use the ENSwR (1–6) theory to offer a way out of this conundrum. Specifically, we deploy a model of population dynamics based on ENSwR as a means to situate the available work on sets of autocatalytic reactions (7, 9, 23–26) within a *bona fide* Darwinian framework. A key conceptual move we make is to lower the level at which units of selection are to be found: while the autocatalytic sets are usually considered as units of selection (because their autocatalysis is taken to be the minimal form of reproduction capable of sustaining ENS (7, 23)), we argue that their components, i.e., relatively simple catalytic molecules, are units of selection. Since the lower-level molecules are incapable of autocatalysis (which could stand in as a proxy for reproduction or self-replication), our niche construction-based approach, built upon a sound articulation of ENSwR, demonstrates how Darwinian evolution can operate prior to the origin of reproducing entities.

Note that throughout this paper, the term “reproduction” refers to a process of multiplication of entities in which parents have a privileged causal input on the traits of their offspring, leading to the presence of vertical lineages linking successive generations (27, 28). We use the term “multiplication” to refer to any process through which entities are being produced; by itself, multiplication entails nothing regarding similarity between producer and products, or among products. “Replication” refers to a special case of reproduction, with high fidelity copying of the entities involved (28–31). This means that reproduction is necessary for ENS, in the classical framework, while replication is facultative (13, 27). “Re-production” refers to cases in which entities are produced recurrently without the privileged causal input of similar entities (parents) (4).

The paper is structured as follows. First, we review the important conceptual aspects of the theory of ENSwR. Second, we describe a model generalizing ENSwR, extracting insights from simulations under a variety of conditions. A key concept we use to structure our model is niche construction, a process tied to a broadened understanding of inheritance mechanisms (32). We show that niche construction can sustain Darwinian evolution even in the absence of reproduction. We suggest that simple chemical catalysts sustain ENSwR when they alter their chemical environment in order to bias future production of entities, thereby increasing their ratio in the population. Finally, we discuss how our model offers a novel theoretical framework that supports ACS-based approaches to the origins of life by addressing some of their theoretical weaknesses.

## 2. Evolution by natural selection without lineage formation

Standard approaches define evolution as changes in the distribution of heritable variation within a population across time. Darwinian evolution is a more specific process in which changes in the distribution of variation are guided by selective pressures, i.e., the result of environmental interactions between members of a population and their environment (5, 12, 13, 27, 33, 34). The environment is seldom defined with ecological precision by evolutionary biology. Rather, it usually refers to the setting that provides selective pressures in which a population or lineage evolves over long timescales (keeping in mind that the environment itself is an evolving system (35)). Standard accounts hold that ENS occurs in any population where there is variation related to fitness and *heredity*. Heredity is held to necessitate reproduction, a causal parent-offspring relationship, which ensures that offspring resemble their parent(s) more than they do other reproducing units in the population (13, 27, 36).

In contrast (see Table 1 for a systematic comparison between standard ENS and ENSwR), the theory of ENSwR suggests that Darwinian evolution occurs in any population where there is variation related to fitness and *memory*. While *heredity* results from mechanisms that ensure the vertical transmission of traits of units of selection, *memory* may result from a more inclusive set of mechanisms that stabilize the distribution of variation *at the level of populations*. Memory can be and often is sustained by the capacity of units of selection to reproduce (reproduction leads to a certain degree of stability at the population level), but memory and reproduction remain different concepts as memory can be outsourced and realized without vertical transmission of traits. The theory of ENSwR hinges on that distinction: memory can be present even in cases where the units of selection fail to reproduce. Niche construction, especially as modeled in section 3, provides empirical grounds and a proof of concept for this claim.

**Table 1.** Differences and similarities between standard ENS and ENSwR.

| Alternative understandings of Darwinian evolution | Standard ENS (13, 27, 30, 49) | ENS<sub>w</sub>R (2, 3, 5) |
| --- | --- | --- |
| Is reproduction necessary? | Yes | No, but the population must be regenerated if new traits are to be explained using ENS<sub>w</sub>R. |
| Darwinian evolution requires *some* stability in the distribution of variation across time. What sustains this stability? | Heredity, i.e., mechanisms of vertical transmission of traits of units of selection. | Memory, i.e., any mechanism that ensures some degree of stability in the distribution of variation in the population (whether through vertical transmission of traits or not). |
| How is fitness defined? (See Box 1 for more details) | Often (but not always) reduced to the reproductive output of a unit of selection; may also include persistence or viability. | A measure of a unit of selection's capacity to increase the ratio of units of selection similar to it in the population. |
| In both cases, Darwinian evolution requires... | • Variation related to differential fitness; fitness entails environmental interactions (natural selection)<br>• Sufficient population size to avoid drift<br>• Some degree of stability in the distribution of variation<br>• Population regeneration to allow small variation to accumulate (required only for origin explanations (27)) |  |

Niche construction has been conceptualized as a process of cross-generational transmission of fitness enhancing effects realized through stable alterations made to the environment by units of selection in past generations (32, 37–41). Beavers, for example, build dams that profoundly alter their ecosystem. Alongside any traits directly transmitted to offspring through genetic inheritance, niche construction by the beaver supports the fitness of generations to come that inherit an environment altered in a beneficial way. Beaver offspring therefore interact with the results of fitness-increasing activities of their ancestors.

Canonical niche construction theory assumes reproduction and builds around it (42): the beavers reproduce, and their niche constructing behavior is evolutionarily important *alongside* this reproduction. In contrast, we believe that niche construction *per se*, i.e., the process of altering the environment in a fitness-increasing way, is relevant even in situations where reproduction is absent. Most importantly, it can accompany other forms of *multiplication* of units of selection, especially in cases where the environment is the main cause of their multiplication. In this perspective, we show that units of selection can alter their environment (niche construction), leading to the re-production *by the environment* of additional similar units of selection in a way that satisfies the condition for Darwinian evolution. Niche construction occurs whether reproduction (multiplication *with* heredity) or re-reproduction (memory-sustaining multiplication without heredity) takes place.

This echoes central insights from the literature on autocatalytic sets of chemical reactions (7, 23, 43). There are many examples of chemical species that, given the right conditions, can affect their environment in this way, often by catalyzing the production of other chemical entities that end up increasing the rate of production of the first species. In hydrothermal vents, where ferrous inorganic barriers separating hydrothermal fluids from ocean water are likely to form (44), a positive feedback loop between CO_2_ fixating reactions and chemical species production occurs, thereby increasing the density of chemical species (45, 46). From the perspective of molecules (e.g., acetate or pyruvate (8)) involved in the process, this is a phenomenon of niche construction: by transforming its environment (i.e., by contributing to a feedback loop), the molecules ensure the proliferation of their type (i.e., increase in density (47)) despite being incapable of reproduction (i.e., these chemical species cannot form lineages).

This phenomenon bridges the gap between evolutionary theorization and the origins of life literature, where ACS-based approaches (and other metabolism-driven ones) are acquiring significant support but struggle to account for the emergence of Darwinian evolution (despite important breakthroughs (48); see section 4 for more details). In the next section, we model dynamics of ENSwR with niche construction. Among other things, this helps explain how reproduction, which is itself an evolved trait (2, 23, 29), may have emerged at the origins of life.

## 3. A model of ENSwR driven by niche construction

The theory of ENSwR has sound conceptual bases but lacks a proof of concept. We designed a model to fill this gap. We built an agent-based model to show how niche construction can sustain Darwinian evolution even in the absence of reproduction, as is the case for chemical species. We then extended the model in stages to show that strong niche construction by chemical species can overcome differential persistence of competitors. In the final version of the model, we add diffusion of units of selection between microenvironments. This provides a foundational inquiry into the possibility that constructed niches are precursors to entities capable of reproduction. More details regarding the model can be found in the Appendix.

### 3.1 The basic model

Our model features a spatial grid of 10 by 10 microenvironments (although simulations presented in sections 3.2 and 3.3 are focused on dynamics occurring within a single microenvironment). These microenvironments represent cell-like pores of alkaline hydrothermal vents (7, 50) populated by various chemical species (where simpler molecules and various gradients lead to synthesis of more complex molecules; see section 4). The chemical species we track, in this model, have different properties related to the (i) rate of production by the local microenvironment, (ii) decay rate, (iii) capacity for chemical modification of the environment, and (iv) diffusion rate. In our simulations, we track only two chemical species, denoted A and B, but we postulate that more are present in the environment. Individuals of both species in a given microenvironment form a single (physically bounded) population of chemical substances.

At regular time intervals all A’s and B’s are terminated. These “extinction” events, which take up a timestep in our simulations, represent occasions when regional environmental conditions exceed the physical tolerance of A and B. These extinction events are akin to paleosterilization events (51). Periodic flows of hydrothermal fluids that exceed the temperature that can be tolerated by local entities are an example of paleosterilization (52) (in the case of prebiotic molecules, the term *paleoextinction* would be better suited than *paleosterilization*). Accordingly, extinction of all A and B units serves two purposes with our model: (i) it mimics the profound instability expected of origin-of-life settings, and (ii) it blocks the material distribution of A and B from directly affecting the future distribution of variation within microenvironments, leaving only their imprint on the environment to affect future distribution of variation. This second point strengthens the case for ENSwR, making it impossible to reduce the dynamics at work to reproduction with vertical transmission. Because of extinction events, long-run dynamics are affected by both the short-term chemical population dynamics that unfold between extinction events, and the degree to which niche construction affects the distribution of variation over longer time scales (across extinction events).

Indeed, the central idea that drives our model is that A and B have the potential to chemically interact with their environment to bias future production of A and B units locally. Each chemical species carries a fixed production bias, and each species is also attributed a value that translates its niche construction capacity. When an individual is introduced in a microenvironment, its niche construction value is added to that microenvironment. Future production of units in that environment is a function of niche construction effects and production bias. In other words, when units of niche constructing species are added to a microenvironment, they increase the chances that more units of the same type are produced in the future. The additive effect of chemical niche construction lasts beyond the periodical extinction of A and B. This operationalizes the notion of niche construction without reproduction as described in the previous section.

We developed two versions of the model: one with unlimited chemical resources available for production and the other with a fixed limit on those resources. The main difference between limited and unlimited resources versions of the model is that in the latter, the production rates of chemical species are independent of one another. In the version where chemical resources are limited, increased production of A’s means a decreased production of B’s (and vice versa) because their production depends on the shared pool of exogenous substrate molecules (“food”). In section 3.2, we compare cases with both limited and unlimited resources. In sections 3.3 and 3.4, we focus on scenarios with limited resources to explore the interplay between niche construction, differential persistence and diffusion across a metapopulation.

Finally, it should be noted that, in this model, time is broken down into units, i.e., timesteps. These timesteps are based on the environmental production rate of units of selection, whereas units of time in many evolutionary models reflect biological reproduction rates instead. For simplicity, we assume that A’s and B’s require the same amount of time between production events, such that we have a unified unit of time. We also simplified the model by assuming that production events are discrete rather than continuous.

### 3.2 Niche-construction VS differential production rates

Our first simulations are meant to establish that chemical niche construction increases the ratio of the niche-constructing units in future generations (see Box 1 on fitness) despite stochastic events such as extinctions (note that we modeled regular extinction events to aid visualization). The objective is to compare the long-term outcome of chemical niche construction to either a high starting production or a mix of high starting production and moderate niche construction (Figure 1). In this design, species A is always better at chemical niche construction than B, and species B starts with a higher production rate (twice that of A in each simulation; note that in the case of scenarios with limited resources, production remains a stochastic process [see Appendix]). We also compared outcomes under unlimited (Figures 1a and 1b) and limited resources (Figures 1c and 1d). Figures 1a and 1c provide results of simulations where A’s have low production rates but chemically modify the local environment in a way that is favourable to their re-production in the long run (A’s niche construction value is 2). Figures 1b and 1d provide results of simulations where B’s also have the capacity to alter the local environment over the long run, but less effectively than A’s (B’s niche construction value is 1). In all cases, A’s come to dominate the population distribution through chemical niche construction.

**Figure 1.**
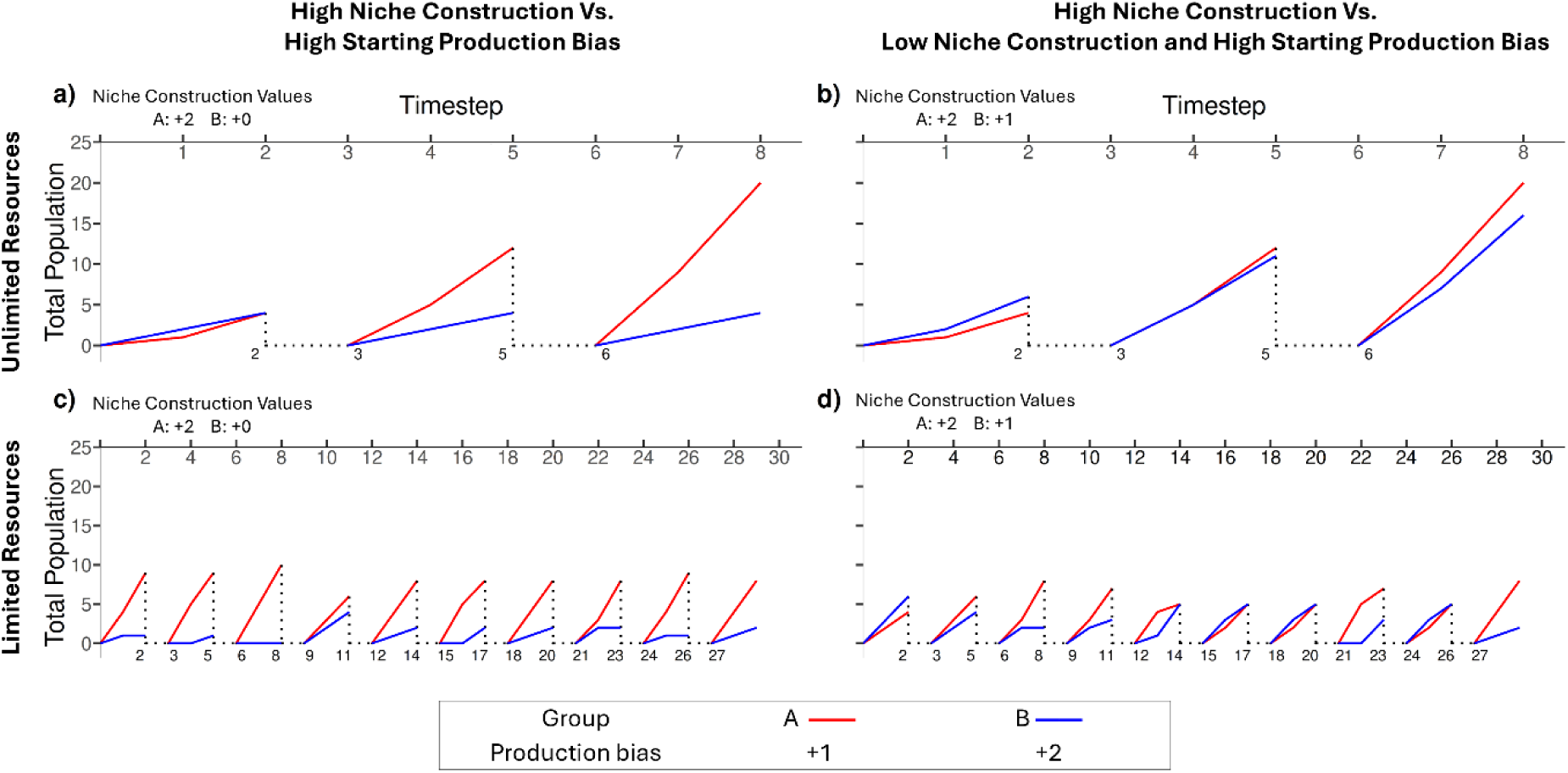
Population dynamics with differential niche construction and differential production biases. This figure illustrates simulations done with two timesteps separating each extinction event (extinction events also count as timesteps), and with the production bias favoring B’s. Extinction events bring the populations down to zero (the dashed lines represent continuity only in the sense that the dynamics happen in the same microenvironment across extinction events). 1a and 1b present scenarios with unlimited resources, while 1c and 1d present scenarios with limited resources. In 1a and 1c, only A’s niche construct. In these situations, A’s come to dominate the population after two extinction events or less. In 1b and 1d, B’s can also niche construct, but A’s do so twice as well as B’s. A’s end up dominating the microenvironment despite starting with lower production rates, but this happens much more slowly.

Some differences must nonetheless be noted. When only A’s niche construct, the relative distribution of A’s in the population reaches 83% when resources are unlimited after only 2 extinction events (the situation is exacerbated when resources are limited). The dynamics are more complex when B’s also niche construct. In both scenarios (limited and unlimited resources), species B initially dominates the population, and under limited resources, it takes more than a dozen extinction events for A’s to reach a frequency comparable to the one they achieve quickly in 1a and 1c (figure 1d: A’s reach 80% of the distribution before the 10^th^ extinction event).

These very basic simulations serve as proof of concept for ENSwR: it shows that change in the relative frequency of A and B results from environmental interactions that feedback as differential multiplication of A’s and B’s within the environment. Recall that A and B do not form lineages (individuals have no privileged causal effect on the generation of specific units of A and B in the future) and undergo periodic “mass extinctions” removing all existing units, entailing complete regeneration of populations. Nevertheless, the dynamics of this simple case, which embodies ENSwR, correspond to Darwinian evolution. Indeed, both ENSwR and the traditional variant, ENS, which entails reproduction and heritability, can be concisely defined as changes in the relative frequency of traits in a population due to natural selection (i.e., selective pressures resulting from environmental interactions) (3, 27, 34). Here, the relative fitness of A and B is expressed as the expected change in the ratio of each in the population, as a result of interactions with the environment. This roughly corresponds to the use of fitness in population genetics (Box 1). However, in traditional population genetics, relative fitness is derived from persistence (or viability) and fecundity, whereas here, relative fitness is a function of production rates and niche construction’s effect on production rates. In the next round of simulations, we add differential persistence to the model. We show that better niche construction can overcome the combination of higher production bias (as in Figure 1) and higher persistence (next section) of competitors.

It could be argued that in situations with limited resources, the effect size of competition for resources between individuals might render the effect of niche construction irrelevant. In our model, this is not the case. First, recall that while production rates are traits of chemical species, this production is mediated by the aggregate effect of niche construction. This aggregate value is a trait of microenvironments, not of units of selection. Second, in figures 1c and 1d there is indeed competition between A and B, at every timestep, to use the limited available resources for molecule production. The effect of competition is strongest at the start because niche construction has not yet impacted the baseline production rate of the microenvironment. Had the results been entirely driven by competition for resources, the species with the highest production bias would always dominate the population distribution. Our results show that an initial advantage in production rates can be overcome by niche construction. Note that results derived from simulations with unlimited resources can only be attributed to the beneficial effect of niche construction on the relative frequency of A (Figures 1a and 1b). Those same processes also determine the outcome when resources are limited, although a greater number of timesteps are needed due to competition for resources modulating the effect-size of niche construction (Figures 1c and 1d). Note also that we use a relatively low quantity of resources in limited-resources scenarios to limit the required computing power. Increased amounts could have an impact on results, likely favoring species A. Having established this proof of concept, the rest of the results are generated under limited resources (where there is an interplay between competition and niche construction), as this better reflects experimental conditions as well as potential *in situ* conditions (7).

### 3.3 Complexifying the model: differential persistence

In the simplest versions of the model (section 3.2), there is no differential persistence (i.e., A’s and B’s have the same decay rate, lasting until the next mass extinction event), such that niche construction and production rates are the only potential features that can lead to change in the distribution of variation. By incorporating differential persistence to the model, an even stronger case is made for niche construction as a way to sustain a response to natural selection in the absence of reproduction. We thus introduced a decay rate for each species, which has no impact on available resources. At each timestep between extinction events, individual A’s and B’s are removed from the population according to their decay rate. To allow decay rates to influence the dynamics of the population, we extended the extinction intervals from 2 timesteps (in the previous section) to 50 timesteps.

First, we provide a null situation where population dynamics are driven only by differential persistence (there is neither niche construction, nor difference in production rates). B’s are the better persistors; A’s decay after only 10 timesteps, while B’s decay after 20. Figure 2a shows that in this situation, B’s dominate the population following each extinction event. Figure 2a illustrates that it takes approximately 20 timesteps to reach predictable equilibrium dynamics after each extinction event. The equilibrium where B’s dominate over A’s is simply a consequence of individual B molecules having a longer residence time due to their lower decay rate as compared to A’s. Note that, in contrast to Figure 1, the baseline production rates for A and B are the same throughout Figure 2. Thus, comparisons with the null case (2a) throughout Figure 2 reveal the interaction between decay rate and niche construction.

**Figure 2.**
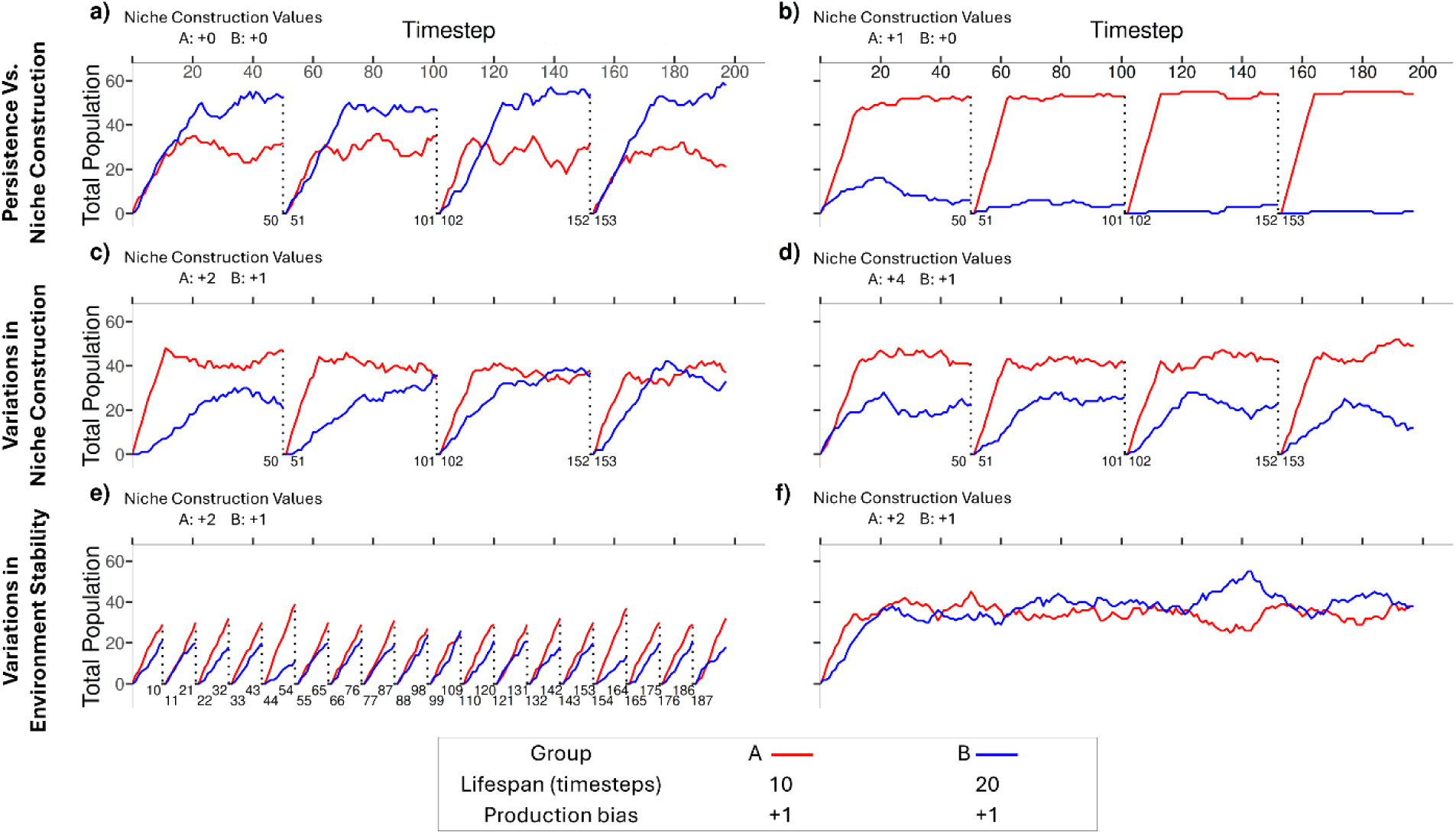
Differential niche construction, differential persistence and extinction-event frequencies. This figure illustrates the dynamics within a single microenvironment in cases where there is differential persistence as well as differential niche construction. Specifically, B’s persist twice as long as A’s throughout the figure. In 2b, A’s niche construction is introduced. In 2c to 2f, A’s and B’s are niche constructors: there is differential persistence *and* differential niche construction. In 2e and 2f, the frequency of extinction events is modulated. All these simulations demonstrate that better niche construction can outcompete better persistence, but that the outcome is dependent on the extent of niche construction or persistence advantages, as well as on specific contextual variables (here, for instance, frequency of extinction events).

In Figure 2b, we re-introduce niche construction. Here, B’s still have better persistence than A’s (20 versus 10 timesteps), but A’s are now capable of niche construction. As expected, A’s very quickly come to dominate the population (as in Figures 1a and 1c). This implies that a niche constructing molecule with the capacity to alter the chemical environment in a way that favours its future production can dominate in a population where competitors are more stable (decay less quickly).

Figures 2c and 2d explore the relationship between persistence and niche construction in cases where both competitors (A and B) are niche constructors. In 2c, B’s survive again twice as long as A’s, but A’s niche construction is twice as efficient. In this situation, the population reaches equilibrium with similar quantities of A’s and B’s present. In this scenario neither A’s niche constructing advantage nor B’s better persistence is enough to make one of the two molecules take over the population. In contrast, 2d illustrates that doubling the niche constructing capacity of A’s causes them to dominate the population even before the first extinction event, with subsequent extinction events increasing the difference. Figures 2c and 2d demonstrate the degree to which better niche constructors dominate a population depends on the interaction with decay rate; stronger niche construction capacity is required when a competitor has a longer residence time due to a low decay rate.

Figures 2e and 2f explore the impact of extinction intervals. The baseline for comparison is scenario 2c. In 2e the extinction events occur on *shorter* intervals (every 10 timesteps), and in 2f there are no extinction events within the interval of the simulation (200 timesteps). In 2e extinction events happen prior to the system reaching equilibrium, so the long-term averages in this system differ from the equilibrium dynamics revealed in 2c. Specifically, in 2e, A’s have a higher prevalence than B’s, while 2c has them on par. This is because the niche constructing capacity of A’s gives them a transient advantage within the first 10 timesteps following an extinction event, and in scenario 2e the extinction events are occurring every 10 timesteps, i.e., before B’s longer decay rate can make a difference. In 2f, we removed extinction events. This simply extends the equilibrium dynamics observed between extinction events (in 2c) throughout the entire simulation. It is easier to see in 2f how there are transient periods where A’s and B’s alternate as the dominant chemical species (due to the stochasticity of production).

### 3.4 Niche construction and diffusion

For this last series of simulations, we added diffusion across a metapopulation (something akin to migration (53)) in order to evaluate the potential of niche constructors to spread beyond their local microenvironment (imagine a network of rocky pores, or compartments on a hydrothermal vent, between which local diffusion of chemical species is possible at a low rate). While chemical diffusion has been used as a basis to understand movement of organisms within a population before (54), here we use the word “diffusion” simply to account for the partially randomized movement of molecules from a microenvironment to neighbouring ones.

The simulations were conducted using a 10 by 10 grid of microenvironments, with border cells linked (so that there is no inherent disadvantage to being produced in a corner microenvironment). At the start of simulations, only one microenvironment produces A’s (this is meant to be a worst-case scenario for A’s and their capacity to spread across the grid). As before, both chemical species must compete for limited chemical substrates. During each extinction event, occurring every 20 timesteps, individual molecules of both types have 1% chance to diffuse from a microenvironment to a *neighbouring* one instead of being completely removed from the metapopulation system. If a niche constructor diffuses to an adjacent microenvironment, its additive effect on production rates (i.e., its niche construction value) is applied to the new environment (without removing its effect on its original microenvironment).

In the scenario depicted in Figure 3, A’s are niche constructors whereas B’s lack the capacity to niche construct. However, B’s have better persistence (20 timesteps) than A’s (10 timesteps). As a result, despite A’s only being produced in a single microenvironment at the start of the simulation, they come to dominate the distribution of variation across the metapopulation after 20 extinction events, and they reach near fixation after 35 (note that we describe the dynamics using extinction events, as the timescales involved are much longer than those relevant in the scenarios depicted by figures 1 and 2). This result is consistent with our previous finding that niche constructors can dominate competitors that persist longer, while extending the result to include the capacity to dominate across a metapopulation when diffusion is possible.

**Figure 3.**
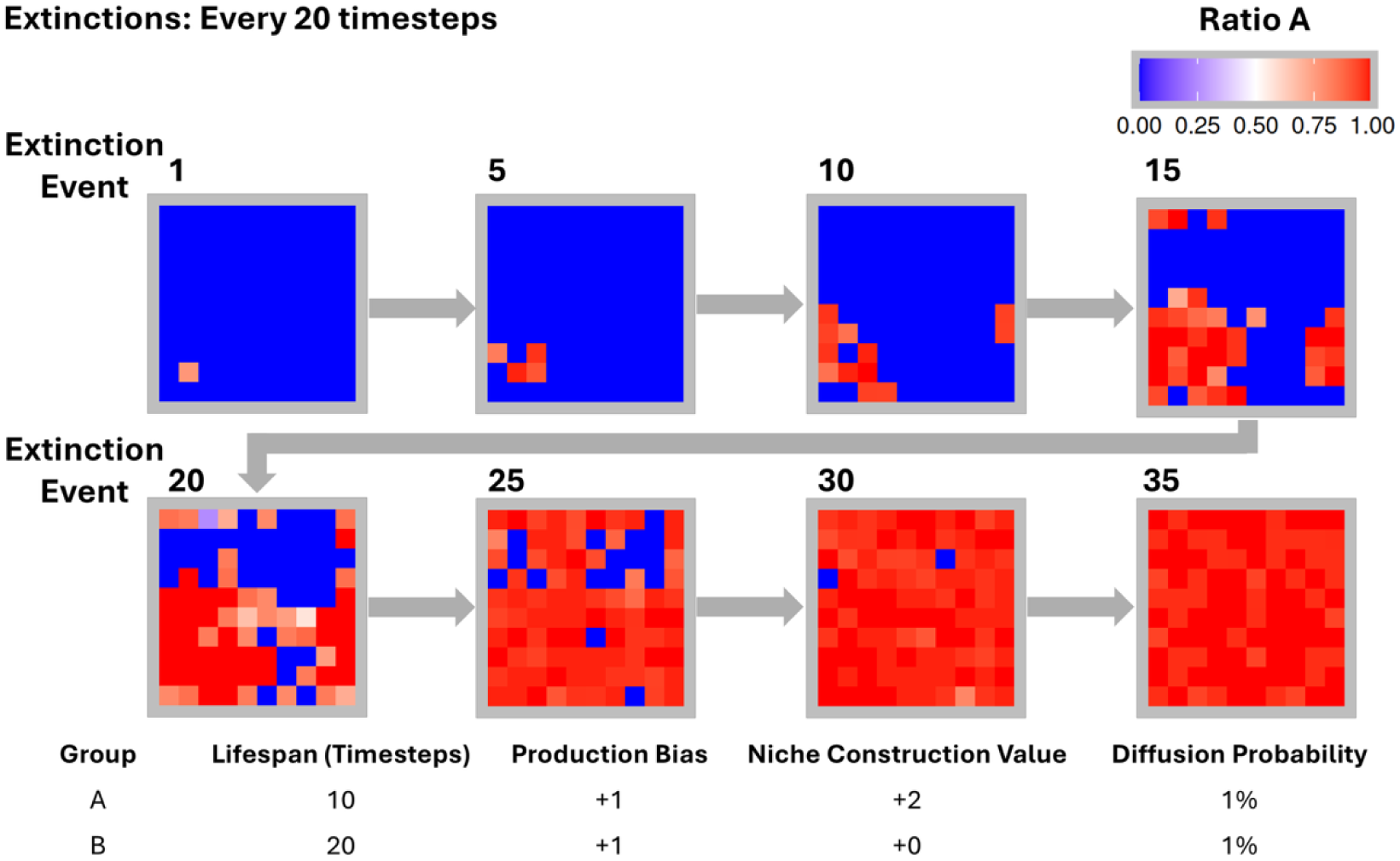
Spread of niche constructors across a metapopulation through diffusion. This simulation introduces the diffusion of units across microenvironments. During each extinction event, individual molecules have 1% chance of being sent from one microenvironment to a neighbouring one instead of being removed from the metapopulation. When this happens, the niche construction effect of the molecule affects its new microenvironment. In this figure and the next, the ratio of A’s and B’s represented by the heat maps are those observed during the last timestep before the indicated extinction event. Species A and B have the same production bias, but only A’s are niche constructors. As a result, even though A’s decay twice as fast as B’s, they spread through the metapopulation quickly.

In figure 4, B’s still persist better, but they are also able to niche construct (B’s niche construction value is 1). Their effect on the environment, however, is weaker than that of A’s (A’s niche construction value is 2). As in Figure 3, A’s are still produced in just a single microenvironment and both types can diffuse to neighboring microenvironments. In this scenario it takes much longer for A’s to spread across the metapopulation and the A’s never reach fixation (after 100,000 extinction events, A’s slightly dominate). The length of this simulation as compared to the others allow us to conclude that this situation leads to an equilibrium between the better niche constructor and the better persistor (as in Figure 2c). As expected, metapopulation dynamics will depend on the strength of the interaction between niche construction and persistence, and fixation only happens when the effect size of niche construction is sufficient to overwhelm that of other traits such as persistence.

**Figure 4.**
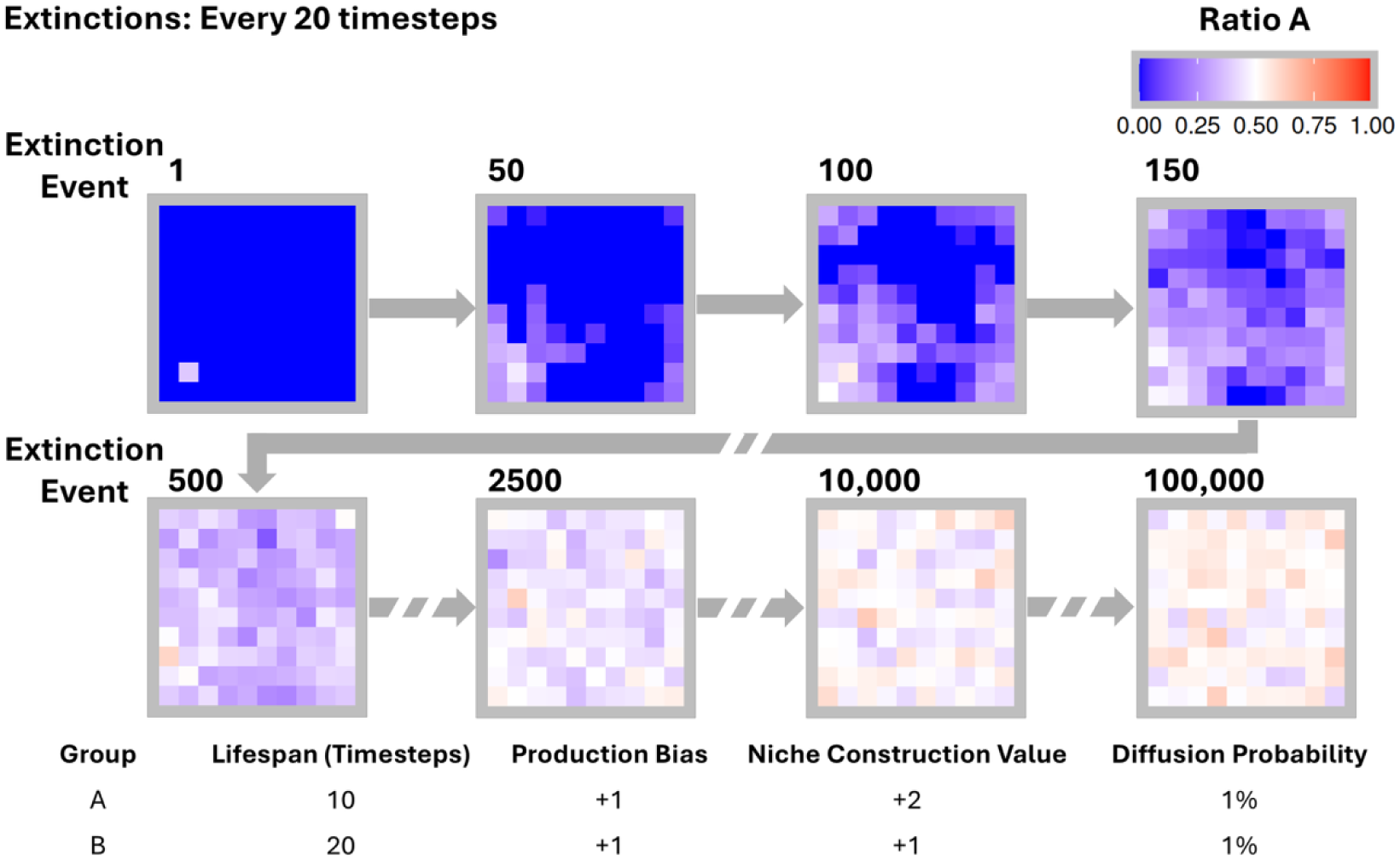
Diffusion, differential niche construction, and differential persistence. This simulation is configured like the one illustrated in Figure 3. The only major configuration difference is the value of B regarding niche construction. While in Figure 3, it did not niche construct, here it does, albeit with a weaker niche construction value than A. B’s longer lifespan partly counteracts A’s better niche construction, such that the metapopulation reaches a near-equilibrium state, with approximately 50% representativity for each species.

## 4. Evolution without reproduction at the origins of life: strengthening ACS-based approaches

The evolutionary model we presented in the above section suggests two important insights for understanding the origins of life: 1- Darwinian evolution can occur without reproduction as long as there is variation related to *memory* (hence the theory of ENSwR); 2- niche construction can drive ENSwR, with memory being the joint effect of particle production by the environment and of their persistence (i.e., their decay rate). In what follows, we wish to extend these insights to empirical case studies. We argue that ENSwR strengthens approaches to the origins of life in which metabolism plays a central role.

Recent enthusiasm for such approaches rests in part on the development, in the past decades, of sound theoretical and experimental breakthroughs surrounding the notion of autocatalytic networks (43, 55). ACSs (also called *reflexively autocatalytic and food-generated sets* or RAFs (56, 57)) is a “set (or cycle) of reactions such that the catalyst of the reactions is synthesized from the food molecules by a series of reactions in the set” (23). A central idea advocated by many researchers in origins of life literature is that ACSs can be units of selection and can form populations that evolve by natural selection. In the first paper where autocatalytic theory was presented (albeit embryonically), Kauffman made explicit that the potential of ACSs to self-replicate is what makes ACSs important to understand early evolution despite their components (i.e., individual molecules) being unable to self-replicate (58). This seminal formulation was refined 15 years later (59). Since then, ACSs have been conceived as the simplest entities capable of reproduction, and therefore of driving ENS according to the classical framework. This established a very specific framework for the origins of life (an alternative to the RNA world hypothesis that then dominated the landscape): if ACSs can emerge from prebiotic chemistry, then they could be at the origin of the first metabolisms, which could have evolved into the first protocells, in which the first templated replication of genetic molecules could have emerged and stabilized more easily. According to this framework, metabolism (ACSs) *precedes* genetic material.

The biggest part of the research regarding ACSs, at least until the turn of the 21^st^ century, was focused on attempting to establish the mathematical and experimental possibility of ACSs arising from prebiotic chemistry (55–57, 60, 61). In the past two decades, however, there has been a sustained interest in the study of ACSs in various empirical settings, such as microbial metabolisms (9, 62), ecosystems (26, 63, 64), and economic systems (65–67). This also includes an interest in empirical prebiotic evolutionary scenarios that gave geochemistry a central place in grounding ACSs approaches to the origins of life (68–70). In what follows, we discuss three challenges that metabolic-first approaches face because of their vexing relationship to evolutionary theory. Adopting ENSwR alleviates these challenges in a way that strengthens frameworks in which metabolism and ACSs feature prominently.

First, ENSwR makes it clear that prebiotic chemistry could have evolved by natural selection even in the absence of templated molecular replication or cell division. This means that theories such as the hydrothermal vent theory (Box 2) have an even better match to available evolutionary theories than researchers in the field have to this day acknowledged. For instance, Williamson recently suggested that Darwinian evolution is incompatible with prebiotic chemistry, and that a new evolutionary mechanism is therefore needed (47). The feeling that prebiotic chemistry requires an alternative theory to ENS is widespread (10, 71–74) but leans heavily on the belief that Darwinian evolution requires reproduction (22). Our simulations show that Williamson’s insights regarding chemical entities indirectly increasing their presence in a population actually sits well with the generalization of Darwinian evolution that the theory of ENSwR makes possible. Indeed, what Williamson emphasizes is the phenomenon of niche construction we theorized above, although he presents it as incompatible with Darwinian evolution. Our models demonstrate how niche construction can drive Darwinian evolution in the absence of reproduction, as is the case with chemical evolution.

Second, past models of ACS evolution have confused evolution (change in the distribution of variation) with Darwinian evolution (change in the distribution of variation that is due to environmental interactions between members of the population and their environment). For instance, Hordijk *et al.* (7) simulated the population dynamics of ACSs (they use the alternative expression RAF) in a model with an architecture similar to ours, in that it features compartments in which various dynamics transform the local chemistry. Whereas we track the dynamics of specific chemical species, Hordijk *et al.* focus on the emergence of specific ACSs. What they show is that there can be changes in the distribution of variation of ACSs across the grid. This is coherent with our results shown in Figures 3 and 4 (section 3.4).

In their model, however, it is unclear in what sense this change in the distribution of variation of ACSs is to be explained by natural selection, i.e., by environmental interactions. Precisely, there is no multiplication of ACSs, only internal transformation due to the production of new chemical species, and it is unclear how ACSs act to maintain or increase the ratio of similar entities in the population. In their model, once the local chemistry activates (catalysts have a small chance of being produced spontaneously but must reach a density threshold to spark local feedback loops), ACSs appear in each compartment (ACSs are produced) but they have no way of interacting with their environment to increase the production of similar ACSs (if anything, they can *hamper* that production by retaining “food” molecules, the constraining resource in their model). There is no selection process, and no way to use fitness to explain population-level changes that occur. Any change in the distribution of variation is due to the internal chemistry that makes ACSs arise and change, a process that is more akin to development than to ENS or ENSwR.

In contrast, in our simulations (especially in Figure 3), the grid goes from being dominated by B’s to being dominated by A’s (and their respective local chemistry) thanks to diffusion *and to A’s more efficient interactions with their environment*. A’s are fitter than B’s, and this explains why they dominate evolutionary dynamics (when they do). In Figure 4, A’s and B’s are approximately as fit, which explains the equilibrium obtained. This showcases why our approach is better equipped to explain how ACSs-based explanations of early prebiotic chemistry correspond to Darwinian evolution. Our crucial move is to recenter evolutionary explanations one level down: the units of selection are neither reproducing protocells, nor ACSs that could give rise to them; they are molecules with catalytic properties that generate chain reactions, which end up increasing their density in a given space due to their interaction with the environment (10, 14, 47).

Third, our approach helps understand the transition between prebiotic chemistry and phenomena driven by classical ENS by offering a novel but preliminary explanation to the origins of reproduction. Indeed, this transition is challenging because, contrary to what is often assumed in the literature, ACSs do not actually *reproduce* when they simply maintain themselves. Reproduction involves, by definition, the *multiplication* of entities. In the way they are traditionally modeled, ACSs maintain themselves and grow (i.e., their components increase in numbers) if the right conditions are met. This phenomenon is loosely tied to self-reproduction (8, 23, 45, 46, 59). This, however, trumps the relationship between reproduction and ENS, two notions that come together only through fitness. Reproduction matters, when it drives evolutionary dynamics, *because* it is a proxy or component of fitness, and being fit means increasing one’s ratio in the distribution of variation within a population. If self-reproduction amounts to maintaining one’s self, then it is more like persistence than like reproduction as usually construed (27, 28, 75).

This matters to ACSs-based approaches to the origins of life because the involved accounts of the transition from prebiotic chemistry to Darwinian evolution assume that metabolic growth leads to reproduction. This is especially relevant to the hydrothermal vents theory at the origins of life (Box 2). In a recent review, Harrison *et al.* (8) suggested that, in hydrothermal vents, ENS could pick up once protocells have developed strong enough positive feedback loops between CO_2_ fixation processes and the production of nucleotides, fatty acids, and other building blocks, because ACS growth would then lead to protocell division. Hence, not only do they assume that reproduction is necessary for Darwinian evolution and that the first units of selection capable of reproduction must have been complex ACSs (protocells), but they also go as far as stating that growth *will* lead to cell division (something that, in certain conditions, does work). In the context of hydrothermal vents and in the very model that they develop, however, protocells are *constrained and dependent on inorganic membranes*. Yet Harrison *et al.*’s paper offers no discussion of how protocells could, in order to undergo division, free themselves of the physical barrier that they relied upon to grow. Their rationale suggests that the result of an evolutionary process, i.e., the emergence of protocell-level reproduction, can be taken for granted. ENSwR can help address this blind spot.

Indeed, our simulations that include diffusion offer a theoretically grounded way to bridge the gap between membrane-bound ACS development and protocell reproduction. The diffusion across inorganic membranes required to sustain the chemical disequilibrium that drives prebiotic processes (76, 77), is the same process that would allow small niche-constructing catalyst molecules to spread to neighboring compartments in a hydrothermal vent. Their spread across microenvironments could therefore lead to the multiplication of the ACSs they give rise too, as they re-produce the ACSs in their new microenvironment through their catalytic activities. This is what we modelled in Figures 3 and 4, although the ACSs are black-boxed in the microenvironment’s weighted production parameter. Once ACSs can be thus multiplied, there would be adequate conditions for invoking multilevel selection theory (with evolutionary dynamics taking place both at the level of molecules and that of ACSs) (30, 78). This all means that ENSwR provides the necessary framework to situate the diffusion of niche constructing molecules as an evolutionary precursor to ACS re-production, and eventually reproduction.

But all this also gives a different outlook on the first protocells and, by extension, on the biological entities that followed their evolution. While evolutionary dynamics were clearly altered once the first cells started reproducing with heredity (79), the early-life equivalent of what are now cells could have been niches being constructed by chemical species. We believe that before being units of selection in their own right, precursors to cells were already something like ecosystems, bridging the gap between the expanding reach of the ACS literature (26, 63, 65) and hydrothermal vents theory. Moreover, this may warrant conceiving of cells and organisms, to this day, as being as much niches (or evosystems (35)) as they are “individuals” or units of selection.

## 5. Concluding remarks: generalizing our results and ENSwR

The Darwinian theory of evolution has outstanding explanatory power and provides grounds for exploring the history and dynamics of biological evolution without focusing on lineage-forming entities. ENSwR expands the reach of ENS by offering a conceptual apparatus (5), a proof of concept (section 3), and an empirical case study (section 4) supporting a version of Darwin’s theory free of the unwarranted yet pervasive metaphysical assumptions according to which units of selection should be reproducers or replicators. By modeling ENSwR with niche construction, the work achieved in this paper provides bases for exploring previously underappreciated evolutionary phenomena in a way that contributes to linking evolution and ecology theories more thoroughly (80). A variety of biological entities (holobionts, ecosystems, genomes, etc.) as well as unsuspected protobiological (catalyst molecules, microenvironments, etc.) and epibiological (social institutions, corporations, etc.) entities may form populations that evolve by natural selection. Furthermore, our approach strengthens the intuition that environmental scaffolds guiding evolution might be more important to the origins of life than the production of specific molecules (e.g., DNA) (35, 81, 82).

Among the many fields in biology that will be influenced by the maturation of ENSwR, we explored the theory’s impact on origins of life research. Specifically, our model solves some of the theoretical problems faced by origins of life literature. Indeed, we have shown that prebiotic chemistry can sustain ENSwR, and can therefore explain the rise of ACSs and ACS re-production without the need to develop an alternative to the Darwinian framework. Our work bridges the gap between chemical and biological evolution in a way that warrants pushing the Darwinian threshold (79) much further back in time than where it has historically been set.

### Box 1 Fitness and ENSwR

Fitness plays two roles in evolutionary biology. One is conceptual: it captures the notion that some entities fare better than others in a given environment. This conceptual role is crucial to the articulation of evolutionary theory, but it has often been tied explicitly to reproduction (83–86). A remodelled definition is therefore necessary to account for the conceptual role of fitness in the context of ENSwR. The second role of fitness is more practical: it is a measure of the conceptual notion. The goal of the present discussion is to identify a way to measure fitness that is coherent with the theory of ENSwR. Indeed, because there is neither reproduction nor lineage formation in our model, we need a more general way to define fitness to argue against statements such as: “an individual’s fitness is identified with the *number of offspring* it produces.” (27)

In population genetics, the understanding and measurement of fitness requires a standardized statistic suitable for predicting population evolution by natural selection (87–90). Standardization starts with absolute fitness (although it is important to note that population prediction requires relative fitness). Absolute fitness of a “type” weighs *viability* (obtained by dividing the number of individuals of a type *after* selection by the number of individuals *before* selection) according to the fertility of that type (obtained by dividing the number of descendants by the number of parents *after viability selection*). Relative fitness converts absolute fitness into a measure of “performance” relative to the other types in the population. Relative fitness is obtained by normalizing absolute fitness in some way; the most common approach is to divide by the highest value of absolute fitness. Relative fitness is necessary because it is not a linear function of absolute fitness, and relative fitness (not absolute fitness) is required for predicting the change of a population due to natural selection. Because relative fitness is a generalized and unitless measure of the performance of a “type” over time, it carries the same meaning when used in the Price equation and population genetics (29, 91, 92).

To develop an account of fitness coherent with ENSwR, we suggest redefining a unit’s fitness as its capacity to influence the ratio of entities with the same or similar trait in the population, which must result from its interaction with the environment. Other ways to define fitness are special cases of this more general formulation. Indeed, reproduction is a specific way by which a unit can increase the ratio of entities with similar traits in the population, but other alternatives also match our definition. For instance, the effect of niche construction on the relative frequency of a type could be weighed in the above measurement procedure just like fertility is; it could even replace fertility altogether. This would still allow predicting population dynamics based on fitness values. In other words, removing reproduction from the *definition* of fitness provides a more general concept that is coherent with cases where reproduction *is* taken into account, but also with cases where reproduction is absent, as in the model presented in this paper.

### Box 2 The Hydrothermal Vents Theory at the Origins of Life

Hydrothermal vents are seafloor structures, such as the Lost City hydrothermal vents (93), whose compositions and topologies are of special geochemical interest. Namely, the rocky formations present far from equilibrium conditions with various gradients enabling and maintaining organic chemistry, such as pH and redox potential gradients (44, 50, 94, 95). These gradients result from the sustained contact of slightly acidic (Hadean) ocean water and alkaline H_2_ rich hydrothermal fluids (44, 96). The thermodynamic and chemical conditions in such settings can lead to the precipitation of (mainly) FeS and Fe(OH)_2_ (76) that then forms a ferrous layer between the two fluids (an inorganic membrane), thereby maintaining and even strengthening the gradients that gave rise to it (76, 97–99). This leads to the formation of rocky pores (i.e., compartments) filled with alkaline nutrient-rich fluid, where the inorganic membrane has been hypothesized to play the role of a precursor to protocellular membrane (8, 44, 50, 94, 95). Specifically, the various gradients maintained and strengthened by inorganic membranes coupled with the available H_2_ promotes the reduction of CO_2_ and therefore organic chemistry (44, 100, 101).

Inorganic membranes can thus serve as evolutionary scaffolds (81) preceding organic membranes. Indeed, in the setting of hydrothermal vents, it has been shown that fatty acids and alcohols can be produced (102, 103), and that they can lead to the formation of fatty acid vesicles (96, 76, 104). These kinds of processes set the ground for the emergence of cells (and thus potentially of life), but more importantly for our purpose, they also imply that positive feedback loops, which can be represented as ACSs, were in place to increase the density of amino acids and other basic chemical building blocks of life (8, 45, 46).

## APPENDIX

Simulations were performed using an agent-based model with timestep progression programmed in Python 3. This model simulates interactions between two key entities: a chemical species and a physiochemical space divided into localized environments (grid cells) (Figure A1). Different chemical species represent molecules with different biases for production by reactions in the local environment. The continuous process of chemical production is discretized into timesteps within the model. Furthermore, particles of a chemical species can chemically modify their local environment to promote the production of more particles of the same species. As such, local chemical environments possess properties conducive to different chemical species, and local environments can be modified when a chemical species is introduced in that environment. These modifications are implemented in the model through the impact of an individual on the value of “niche property” in its local environment. Periodically, the environment suffers a chemical ‘mass-extinction’ during a timestep. Such an event causes all species of chemical molecules to be removed from the whole environmental space (i.e., flushed or destroyed). However, the localized environments are not physically destroyed in this process, allowing prior chemical modifications of the physical environment to persist until the chemical process of molecule production is re-established in each microenvironment.

**Figure A1.**
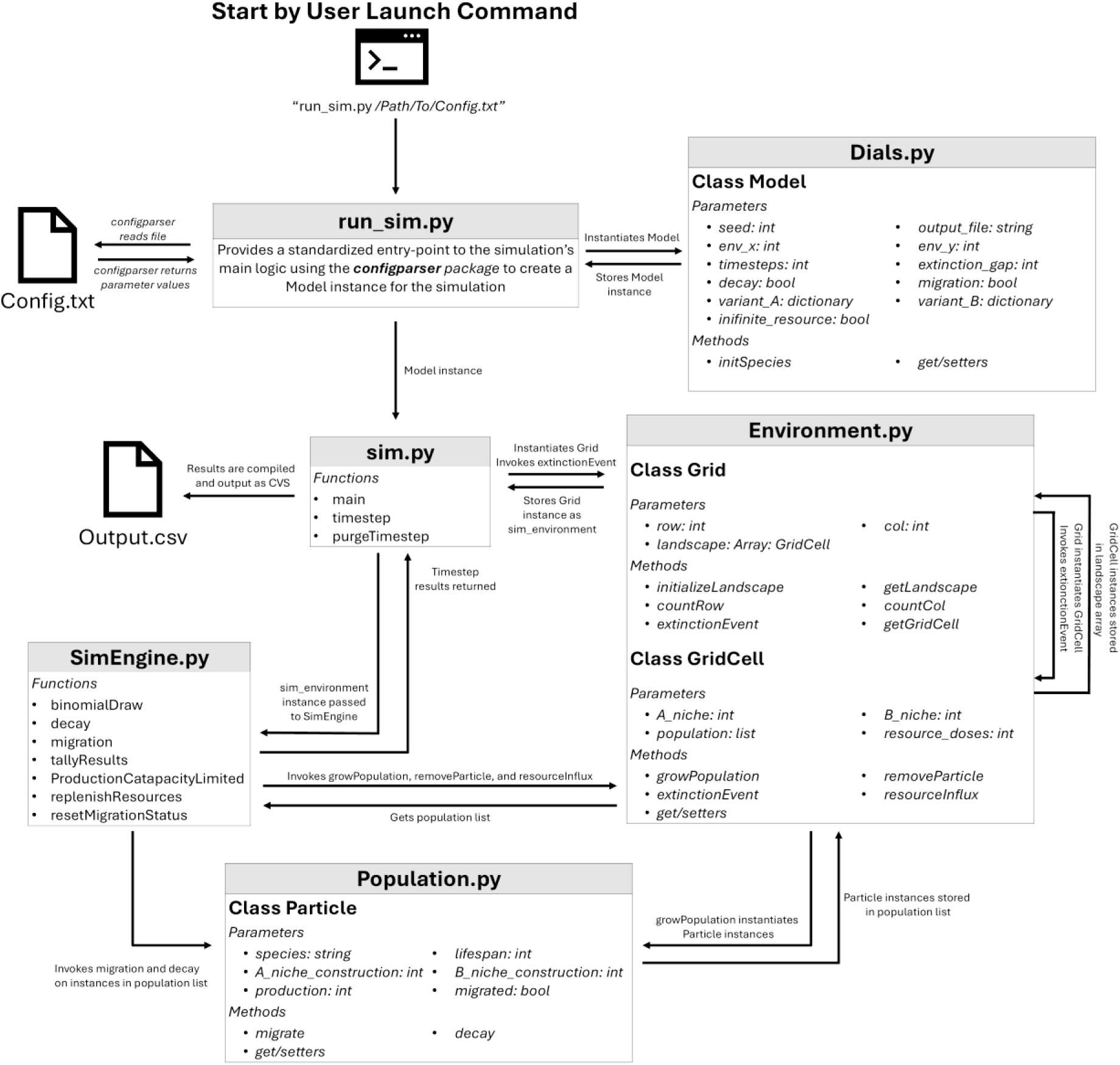
Relationship diagram depicting flow of information through interactions between the modules and classes of the ENSwR simulation to produce output.

This model has independent settings to enable or disable resource limitation, particle decay, and migration conditions. Using combinations of these conditions, we can create eight (2^3^) unique modes of simulation.

## Chemical species production under unlimited resources

When resources are unlimited, creation of the individual particles of a chemical species is determined by the sum of its species-specific production bias and the chemical properties of the environment (Table A1, Equation 1). In this scenario, production of one chemical species does not affect the production of another chemical species, and there are no imposed maximum limits on particle production rate for any of the chemical species.

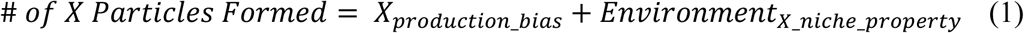

**Table A1.** Description of all parameters comprising a configuration file.

| Name | Data Type | Description |
| --- | --- | --- |
| *[Dials]* |  | Header for model settings |
| seed | int | Optional declaration for random number seed |
| timesteps | int | Number of timesteps to run the simulation |
| extinction_gap | int | Timesteps between mass-extinctions |
| environment_x | int | Size of Environment grid (x dimension) |
| environment_y | int | Size of Environment grid (y dimension) |
| output_file | string | Filepath to direct output file |
| decay | bool | Toggle whether model includes Particle decay |
| migration | bool | Toggle whether to include migration |
| infinite_resources | bool | Toggle whether to include resource limitation |
| *[Type_A/B]* |  | Header for Particle types (A or B) settings |
| species | string | Species of Particle (A or B) |
| A/B_niche_construction | int | Niche construction value for species |
| production_bias | int | Innate production for species |
| lifespan | int | Timesteps to persist in decay models |

## Chemical species production under limited resources

In this scenario, the production of a chemical species is limited by the availability of shared precursor molecules. These are handled generically within the model by a generalized “resources” parameter. Resource utilization is modeled independently within each localized environment. In our simulations, each localized environment replenishes to a fixed number of resource “doses” every timestep, which reflects the baseline chemical production of precursor molecules within the local environment. Each dose of resource is chemically transformed to produce one chemical species of a specified type, such that each localized environment will spontaneously produce a total of five chemical particles during each non-extinction timestep of the model. This process is modeled as a Bernoulli experiment occurring each timestep. In these experiments, consumption of each resource “dose” is treated as a trial where probability of outcomes can be determined by the accumulated effects of a chemical species’ relative rate of baseline production (bias) and the niche-constructed chemical properties of its environment (Equation 2). This imposes resource competition for production of different chemical species.

For each Bernoulli trial at grid cell *n, m*

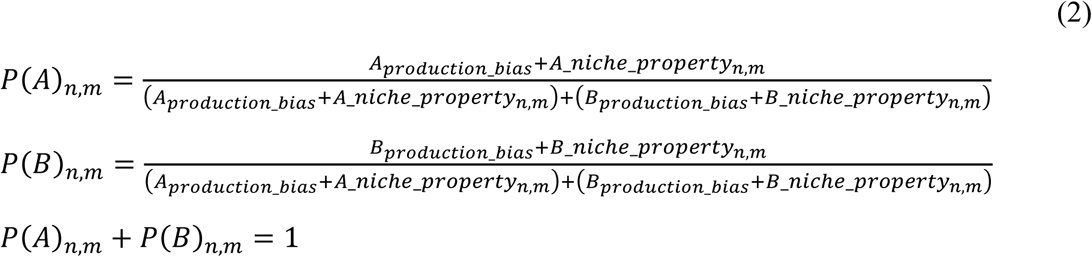

## Particle Decay

Decay reflects the capacity by some molecules to have greater molecular stability than others and therefore to persist longer. In these models, chemical species have a lifespan parameter which specifies how many timesteps a particle of that type can persist. When a particle has exceeded its lifespan, it is considered to have decayed, and it is removed from the environment. Without decay, chemical particles persist until they are destroyed in a mass-extinction timestep.

## Diffusion (an analogue to “migration”)

Chemical diffusion is well understood, and because it is mathematically convenient, it is often used as a model to approximate large scale patterns of animal movement. Here we are looking at something else, i.e., movement of particles caused by a mechanical force in the environment. We call this “particle diffusion” simply to refer to the partially random movement of chemical species. We implement diffusion via a random process of individual particle movement between neighbouring local environments (grid cells). In this model, particle movement is a discrete process coordinated by the SimEngine at discrete time points (*i.e*., we discretized and marginalized the effect of the continuous process). We update particle movement between local environments during each mass-extinction timestep. At this time point, each particle has a 1% chance of surviving the extinction event, but gets relocated to an adjacent localized environment. These new arrivals immediately interact with their new local environment to modify the relevant chemical niche parameter(s). As described above, any modifications made to a local environment’s properties persists within that local environment.

Additionally, for the purposes of demonstrating dynamics in an extremely challenging case, particle species “A” was assigned a positive production bias only within a single local environment. Hence, for species “A” to populate new local environments, individual A’s must “diffuse” to apply their species’ unique modification to the local chemical niche-property. As its production bias value is 0 in any new environment, all future production of “A” in a new environment is determined by the effect of its niche construction (implemented as the value of its “niche” parameter). Effectively, without “diffusion” enabled, “A” is restricted to a single local environment within the grid.

## Running the Model

The model is specified according to user-defined settings in a configuration file. Running a simulation according to the specified model requires calling the *run_*sim module with the pathway for the configuration file as its argument. Parameters in the configuration file are described in Table A1. The *run_sim* module uses the configuration file to construct a model object. The model is then sent to the *sim* module, which creates the environmental grid and begins computing the simulation timesteps. In doing so, the *sim* module (1) invokes a *SimEngine* module responsible for probability calculations, (2) coordinates the environment and particle objects to perform the steps required for the simulation, and (3) logs the data to be sent back to the *sim* module, where it is formatted and written to files after all timesteps are completed (Figure A1).

